# The fully activated open state of KCNQ1 controls the cardiac “fight-or-flight” response

**DOI:** 10.1101/2024.07.02.601749

**Authors:** Panpan Hou, Lu Zhao, Ling Zhong, Jingyi Shi, Hong Zhan Wang, Junyuan Gao, Huilin Liu, Joan Zuckerman, Ira S. Cohen, Jianmin Cui

**Affiliations:** Department of Biomedical Engineering, Center for the Investigation of Membrane Excitability Disorders, Washington University, St. Louis, MO 63130, USA; Department of Physiology and Biophysics, Stony Brook University, and Institute for Molecular Cardiology, Stony Brook, NY 11794, USA

**Author notes:** Send correspondence to; Jianmin Cui, Panpan Hou. Current address: Dr. Neher’s Biophysics Laboratory for Innovative Drug Discovery, State Key Laboratory of Quality Research in Chinese Medicine, Macau University of Science and Technology, Taipa, Macao SAR, China.

**Keywords:** I_Ks_ channel, Phosphorylation, “fight-or-fight” response, Long QT syndrome, anti-arrhythmia

## Abstract

The cardiac KCNQ1+KCNE1 (I_Ks_) channel regulates heart rhythm in both normal and stress conditions. Under stress, the β-adrenergic stimulation elevates the intracellular cAMP level, leading to KCNQ1 phosphorylation by protein kinase A and increased I_Ks_, which shortens action potentials to adapt to accelerated heart rate. An impaired response to the β-adrenergic stimulation due to KCNQ1 mutations is associated with the occurrence of a lethal congenital long QT syndrome (type 1, also known as LQT1). However, the underlying mechanism of β-adrenergic stimulation of I_Ks_ remains unclear, impeding the development of new therapeutics. Here we find that the unique properties of KCNQ1 channel gating with two distinct open states are key to this mechanism. KCNQ1’s fully activated open (AO) state is more sensitive to cAMP than its’ intermediate open (IO) state. By enhancing the AO state occupancy, the small molecules ML277 and C28 are found to effectively enhance the cAMP sensitivity of the KCNQ1 channel, independent of KCNE1 association. This finding of enhancing AO state occupancy leads to a potential novel strategy to rescue the response of I_Ks_ to β-adrenergic stimulation in LQT1 mutants. The success of this approach is demonstrated in cardiac myocytes and also in a high-risk LQT1 mutation. In conclusion the present study not only uncovers the key role of the AO state in I_Ks_ channel phosphorylation, but also provides a new target for anti-arrhythmic strategy.

**Significance statement:** The increase of I_Ks_ potassium currents with adrenalin stimulation is important for “fight-or-flight” responses. Mutations of the IKs channel reducing adrenalin responses are associated with more lethal form of the type-1 long-QT syndrome (LQT). The alpha subunit of the IKs channel, KCNQ1 opens in two distinct open states, the intermediate-open (IO) and activated-open (AO) states, following a two-step voltage sensing domain (VSD) activation process. We found that the AO state, but not the IO state, is responsible for the adrenalin response. Modulators that specifically enhance the AO state occupancy can enhance adrenalin responses of the WT and LQT-associated mutant channels. These results reveal a mechanism of state dependent modulation of ion channels and provide an anti-arrhythmic strategy.

## Introduction

In the heart, the voltage-gated KCNQ1 potassium channel associates with its KCNE1 modulatory subunit to form the slowly activating delayed rectifier I_Ks_ (KCNQ1+KCNE1) channel, which contributes to the repolarization of cardiac action potentials (1). Congenital mutations in both *KCNQ1* and *KCNE1* genes can cause long QT syndrome (LQTS), which shows a prolongation of the QT wave interval in a patient’s electrocardiogram (ECG) and a propensity to ventricular tachyarrhythmias that may lead to cardiac arrest and sudden death (2–5). LQTS is associated with variants of various ion channels and other proteins (6), but Long QT syndrome type 1 (LQT1), is due to loss-of-function mutations in KCNQ1, and is the most common congenital LQTS, accounting for more than one third of all cases (7, 8).

I_Ks_ plays a key role in controlling the heart rhythm, especially under stressful conditions that require increased cardiac output following the β-adrenergic stimulation. This process is also known as the cardiac adaptation (or the “fight-or-flight” response), during which the neurotransmitter norepinephrine (NE) activates the cardiac β-adrenergic receptors, which enhances both inward (e.g. ICa_L,_ I_F_) and outward currents (I_Ks_ and to a much smaller extent I_Kr_). It achieves these changes by increasing the intracellular cyclic adenosine monophosphate (cAMP) concentration. In the case of I_Ks_, protein kinase A (PKA) is then activated and it phosphorylates the I_Ks_ channel (9, 10). The phosphorylation of the channel dramatically increases I_Ks_, thereby promoting the action potential shortening that allows sufficient diastolic refilling during an increase in heart rate (9, 11, 12) (**Fig. 1A,B**). Clinical studies found that LQT1 patients tend to develop cardiac arrest when the β-adrenergic pathway is activated (for example, when they exercise or are under emotional stress) (9, 12, 13), and LQT1 mutations that impair the β-adrenergic (cAMP) regulation of mutant-I_Ks_ channels pose much higher risk of arrhythmia and sudden death to patients than others (14). However, the molecular mechanism underlying the I_Ks_ channel phosphorylation remains elusive, and the lack of means to rescue the defective cAMP response of high-risk variants impedes novel therapeutic options.

**Fig. 1.**
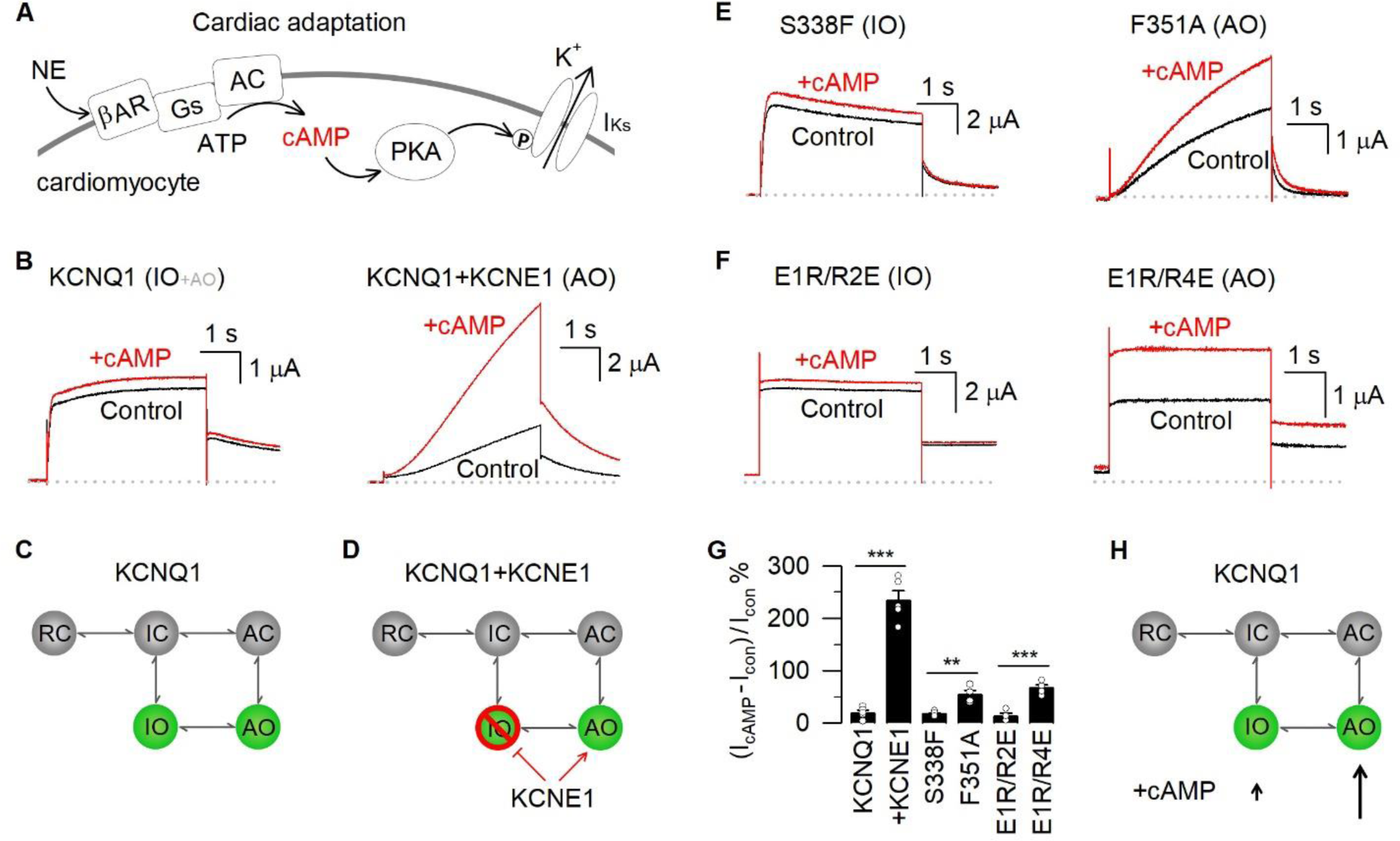
The AO state of KCNQ1 has higher cAMP sensitivity than the IO state. **(A)** Cartoon model to show the β-adrenergic-dependent upregulation of I_Ks_ currents in the heart. Abbreviations: NE, norepinephrine; βAR, β-adrenergic receptor; Gs, G-protein subunits; AC, adenylyl cyclase; PKA, Protein Kinase A; P, phosphate. **(B)** Representative currents of KCNQ1 and KCNQ1+KCNE1 channels, expressed in *Xenopus* oocytes, before (black) and after (red) adding 0.5 mM cAMP. The test pulse was +40 mV for 4 s. **(C, D)** Cartoon scheme to show the gating mechanism of the KCNQ1 channel. The KCNQ1 channel predominantly opens in the IO state, while KCNE1 suppresses the IO state but enhances the AO state. **(E-F)** Representative currents of S338F and E1R/R2E (open in the IO state), and F351A and E1R/R4E (open in the AO state) before (black) and after (red) adding 0.5 mM cAMP. The test voltage was +40 mV for 4s and then stepped back to −40 mV. **(G)** cAMP induced current increases from KCNQ1 alone (19.6 ± 4.8%), KCNQ1+KCNE1 (234.3 ± 18.1%), S338F (17.6 ± 1.9%), F351A (54.1 ± 8.0%), E1R/R2E (13.7 ± 5.1%), and E1R/R4E (66.6 ± 6.4%). Error bars are standard error of the mean (S.E.M.) in this study, n≥4. In all figures *, p < 0.05; **, p < 0.01; and ***, p < 0.001; unpaired Student’s t-test. **(H)** A cartoon scheme to illustrate that the AO state is more sensitive to cAMP than the IO state.

KCNQ1, also known as K_V_7.1, belongs to the classic homo-tetrameric voltage-gated potassium (K_V_) channel subfamily (2, 3, 5). Each subunit contains six transmembrane segments (S1-S6), with the S1-S4 forming the voltage sensing domain (VSD) and the S5-S6 constituting the pore domain (15–18). The VSD activates in response to membrane depolarization in two measurable steps, first to an intermediate (I) state and then to the activated (A) state. Unique to the KCNQ1 channel, the channel pore opens when the VSD is at either I or A state, giving rise to two distinct open states, the intermediate-open (IO) and activated-open (AO) states (18–24). The coupling between VSD activation and pore opening for the IO and AO states follows a dynamic “Hand-and-Elbow” gating process, such that movements of S4 in the VSD and subsequent movements of the S4-S5 linker triggers pore opening by interacting with S6 of the same subunit, and then with S5 and S6 of a neighboring subunit (22). The IO and AO states have different gating and modulation properties. For example: 1) Distinct VSD-pore coupling mechanisms are responsible for the IO and AO states. Multiple LQT1 mutations (e.g. S338F (25)) were found to specifically suppress the VSD-PD coupling of one (AO) state, leaving the mutant channels open only in the other (IO) state (20–22). 2) KCNQ1 predominantly opens in the IO state, while KCNE1 suppresses the IO state but enhances the AO state, so that the I_Ks_ channel opens only in the AO state (19–23) (**Fig. 1C,D**). 3) A small molecule compound ML277 (26) can increase the AO state current by specifically enhancing the VSD-pore coupling, partially mimicking the KCNE1 function (18, 21, 22).

In this study, we show that the above unique properties of gating and modulation of KCNQ1 are the bases for the mechanism of I_Ks_ channel phosphorylation. Our study uncovers the key role of the KCNQ1 AO state in the channel phosphorylation, and based on these findings, we propose that the AO state from native I_Ks_ channels can be a new target to rescue the defective cAMP sensitivity of high-risk LQT1 mutations. First, using mutagenesis tools that selectively favor the IO and AO states, we found that the AO state of KCNQ1 is responsible for the cAMP response. KCNE1 largely enhances the AO state, and therefore boosts the cAMP-induced current increase of KCNQ1. Taking advantage of the unique property that ML277 specifically enhances the AO state, we next found that ML277 can efficiently increase the cAMP sensitivity of both the WT KCNQ1 channel (without KCNE1 association) and the native I_Ks_ channel (with unsaturated KCNE1 association) from guinea pig cardiomyocytes. These results lead to a new strategy for the treatment of lethal LQT1 associated with mutations that impair cAMP sensitivity of I_Ks_, in which pharmacological reagents can be used to enhance the AO state of the mutant I_Ks_, thereby recovering the cAMP sensitivity of the mutant channel. To demonstrate this strategy, we studied a high-risk mutation R243C of KCNQ1, which reduced cAMP sensitivity of the I_Ks_ channel, and found that enhancing the AO state occupancy by ML277 rescues the defective cAMP sensitivity of R243C. Taken together, our findings not only reveal the molecular mechanism of the PKA-dependent I_Ks_ phosphorylation but also provide a promising new anti-arrhythmic strategy, i.e. to enhance the AO state occupancy of mutant KCNQ1 channels, for high-risk LQT1 variants.

## Results

### The AO state of KCNQ1 has a higher cAMP sensitivity than the IO state

It is well established that the KCNQ1 channel shows little sensitivity to β-adrenergic stimulation via the cAMP-dependent protein kinase A, but when KCNQ1 is associated with KCNE1, cAMP induces a large current increase (14, 27–30) (**Fig. 1A,B**). The KCNE1 boosted cAMP effect increases the β-adrenergic activation of I_Ks_ to accelerate the repolarization of cardiac action potentials in adaptation to stress conditions (11, 12, 14, 27–30). However, why the KCNE1 subunit is required for such a cAMP-induced current increase remains unclear. To solve this long-standing puzzle, in this study, we applied membrane permeable cAMP (8-Br-cAMP) to mimic the β-adrenergic stimulation of the KCNQ1 and I_Ks_ channels expressed in *Xenopus* oocytes. We found that 0.5 mM cAMP had minimum effect on the KCNQ1 current amplitude (19.6 ± 4.8% increase) and the conductance-voltage (G–V) relation, but tripled the KCNQ1+KCNE1 current amplitude (234.3 ± 18.1% increase) and shifted the G–V relation to more negative voltages (**Fig. 1B** and **Fig. S1**). These results obtained from *Xenopus* oocytes are consistent with findings in mammalian cells like Chinese hamster ovary (CHO) cells, Itk^−^ mouse fibroblast (LM) cells and human embryonic kidney (HEK) 293 cells (11, 31, 32).

We then investigated the mechanism for the different cAMP sensitivity in KCNQ1 and I_Ks_ channels. It has been shown that, in response to depolarization, the VSD of KCNQ1 undergoes two steps of activation to the intermediate and activated states (19, 20, 22) (I and A, **Fig. 1C,D**). Through a dynamic VSD-pore coupling process, both the I and A state VSD movements induce pore openings, thus the channel has two different open states, IO and AO (18–20, 22) (**Fig. 1C,D**). The two open states show distinct functional properties including time- and voltage-dependence, ion permeation, and drug sensitivity (19–22). The KCNQ1 channel primarily opens to the IO state, while KCNE1 suppresses the IO state but enhances the AO state and thus the I_Ks_ channel exclusively opens to the AO state (19, 20, 22, 23).

Based on these previous results, we hypothesize that the IO and AO states have different sensitivities to the cAMP stimulation and the AO state is more sensitive to cAMP than the IO state, thus the KCNQ1 channel that has primarily the IO state phenotype shows low cAMP sensitivity, while the I_Ks_ channel that has the AO state phenotype shows enhanced cAMP sensitivity. To test this hypothesis, we utilized our previously identified two pairs of mutant channels that specifically open in the IO or AO state and studied their responses to cAMP stimulation. The first pair, S338F (a long Q-T syndrome LQT1-associated mutation (25)) and F351A, selectively disrupt the VSD-pore coupling at the AO and IO state, and thus leave the mutant channels open only in the IO and AO state, respectively (20–22). The second pair, E160R/R231E (E1R/R2E) and E160R/R237E (E1R/R4E),arrest the VSD at the I and A state, and make the mutant channels constitutively open in the IO and AO state, respectively (19, 20, 22).

We found that the application of 0.5 mM cAMP increased the currents of E1R/R4E and F351A that open in the AO state, but enhanced E1R/R2E or S338F that open to the IO state to a lesser extent (**Fig. 1E-G**). These results demonstrate that, consistent with our hypothesis, the AO state is more sensitive to cAMP than the IO state (**Fig. 1H**), which answers the long-standing question of why KCNE1 can boost the cAMP sensitivity of KCNQ1: the association of KCNE1 in I_Ks_ enhances the cAMP-sensitive AO state and suppresses the less cAMP sensitive IO state, and therefore boosts the cAMP-induced current increase of I_Ks_.

### Increasing the AO occupancy by modulators enhances cAMP sensitivity of KCNQ1

The above results not only show the critical role of the AO state in the KCNQ1 channel cAMP sensitivity, but also suggest that the KCNE1 subunit may not be required for the cAMP-dependent modulation process. Based on this mechanism, we expect that a specific increase of the AO state occupancy in KCNQ1 will enhance its cAMP sensitivity.

We have previously studied the functional effects of the small molecule ML277 on KCNQ1 channels and found that it exclusively enhances the AO state occupancy of the channel. ML277 alters the phenotype of the channel by inducing the AO state with increased current amplitude, a slower activation rate, a right-shifted G–V relation, and a decreased Rb^+^/K^+^ permeability ratio (21). All these modulations are consistent with the mechanism that ML277 mimics the KCNE1 effects on KCNQ1 to enhance the AO (**Fig. 2A**). It was demonstrated that ML277 binds to the interface between the VSD and the pore of KCNQ1 (17, 18, 21), and specifically enhances the VSD-pore coupling for the AO state. We therefore tested if ML277 is capable of increasing cAMP sensitivity of KCNQ1 channels similar to KCNE1. Consistent with our previous results, 1 µM ML277 increased the current amplitude of the slow component without affecting the fast component (**Fig. 2B**), indicating that the channels opened with an enhanced AO state occupancy (21, 22). Strikingly, in the presence of 1 µM ML277, the KCNQ1 channels were more sensitive to cAMP, such that cAMP induced larger current increases as compared to KCNQ1 without ML277 (**Fig. 2B-E**). A higher ML277 concentration (3 µM), which induced a higher AO state occupancy, further boosted the cAMP sensitivity of KCNQ1 (**Fig. 2E**). We also compared the voltage-dependent activation and found that ML277 and cAMP significantly increased the total maximal conductance, while the G-V relation after adding ML277 and cAMP showed a shift to more positive voltages of ∼6 mV (**Fig. 2F**). This shift was mainly induced by ML277 due to the increased AO occupancy, which shifted the G-V relation in the direction of the G-V of the AO state as in the I_Ks_ channel (18, 21). These results confirmed the key role of the AO state in modulating the cAMP-dependent phosphorylation of both KCNQ1 and I_Ks_ channels.

**Fig. 2.**
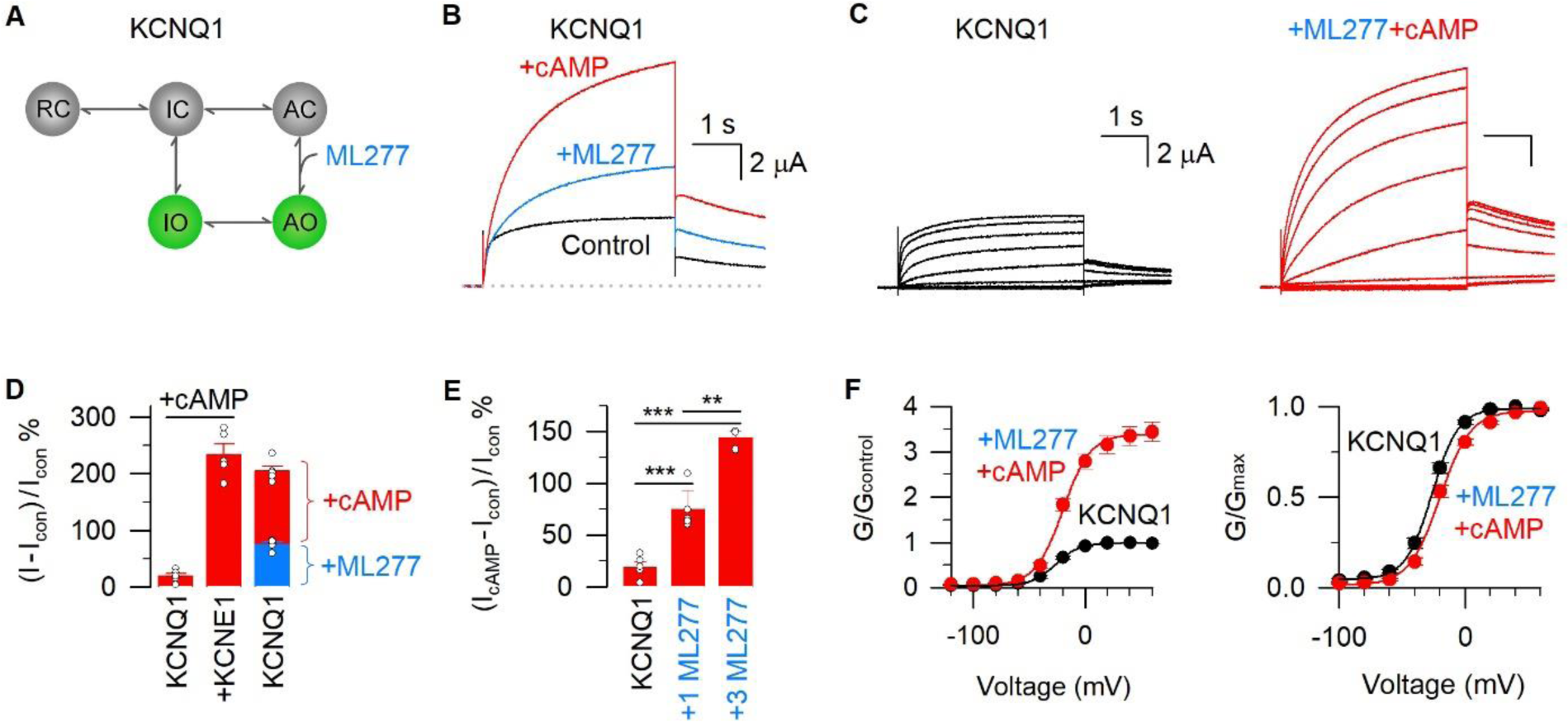
Increasing AO occupancy by ML277 enhances the cAMP sensitivity of KCNQ1. **(A)** Cartoon schemes to illustrate that ML277 binds at the interface between two neighboring KCNQ1 subunits and specifically enhances the AO state of the KCNQ1 channel, mimicking the KCNE1 effects (21). **(B)** Representative currents of KCNQ1 channel before (black) and after (blue) adding ML277, and then after adding cAMP (red). The test voltage was +40 mV for 4s and then stepped back to −40 mV. **(C)** Representative activation currents of KCNQ1 before (black) and after (red) adding ML277 and cAMP. **(D)** Current increases of KCNQ1 after adding ML277 (blue, 75.2 ± 4.0%) and cAMP (red, 205.1 ± 8.5%). The same data for cAMP-induced current increases of KCNQ1 (19.6 ± 4.8%) and KCNQ1+KCNE1 (234.3 ± 18.1%) channels are shown. All n≥4. **(E)** The cAMP-induced current increases of KCNQ1 before (19.6 ± 4.8%) and after adding 1µM (74.9 ± 18.1%) and 3 µM (144.2 ± 5.8%) ML277. All n≥3. **(F)** Left, G-V relations of KCNQ1 before (black) and after (red) adding ML277 and cAMP normalized to control. Right, normalized G-V relations of KCNQ1 before (black) and after (red) adding ML277 and cAMP. All n≥4.

To further validate the predominant role of the AO state for cAMP sensitivity, we then proceeded to investigate whether other compounds with the ability to modulate the AO state can likewise enhance the cAMP sensitivity. C28 is a small molecule I_Ks_ activator that was found to bind to the VSD of KCNQ1 (33). Consistent with our previous studies (33), C28 enhanced I_Ks_ but did not enhance the current of KCNQ1 (**Fig. 3A**). This result was due to C28’s preferential enhancement of the AO state occupancy (33). We found in the current study that 3 µM C28 accelerated the activation and slowed down the deactivation of KCNQ1 currents, while it decreased the current amplitude (**Fig. 3B, C**). These observations are consistent with our previous results (33). Remarkably, after adding C28, 0.5 mM cAMP induced a significantly larger current increase than that of the WT KCNQ1 (105.3 ± 18.3% vs 19.6 ± 4.8%, **Fig. 3B-D**). We also tested the voltage-dependent activation and found that, after adding C28 and cAMP, the G-V relation of KCNQ1 showed an increased total maximal conductance and a ∼35 mV shift to more negative voltages (**Fig. 3E, F**). This G-V shift was part of the activation effect of C28 (33). Together, these results support that C28, which enhances the AO state occupancy by a different mechanism than ML277, can also sensitize the KCNQ1 channel to cAMP.

**Fig. 3.**
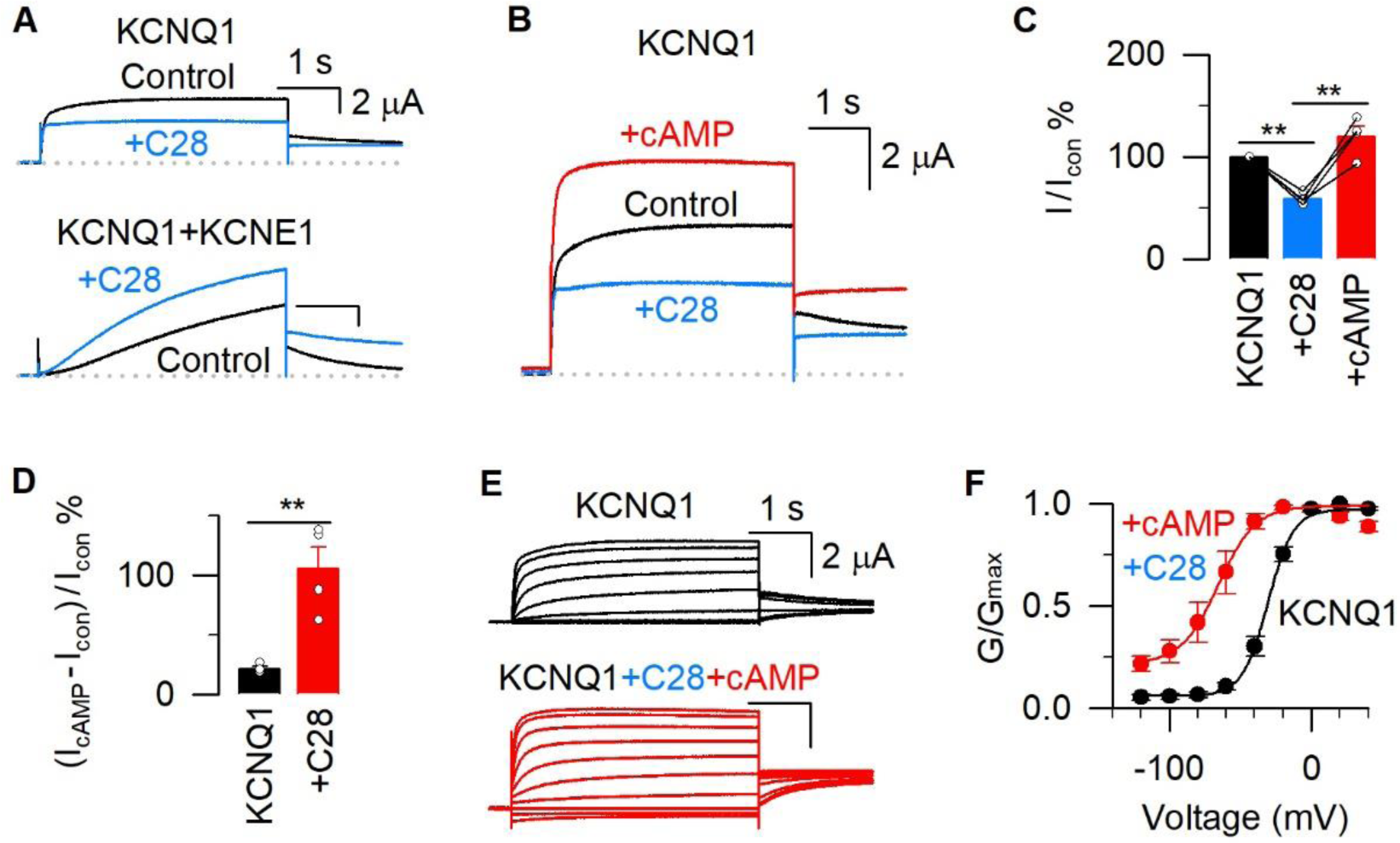
C28 also boosts the cAMP effects of KCNQ1. **(A)** Representative currents of KCNQ1 and KCNQ1+KCNE1 before (black) and after (blue) adding 3 µM C28. **(B)** Representative currents of KCNQ1 channel before (black) and after (blue) adding 3 µM C28, and then after adding cAMP (red). **(C)** Normalized current changes of KCNQ1 after adding C28 (blue, from 100 % to 58.7 ± 2.9%) and then after adding cAMP (red, from 58.7 ± 2.9% to 119.8 ± 9.6%). **(D)** The cAMP-induced current increases of KCNQ1 (19.6 ± 4.8%) and KCNQ1 after adding C28 (105.3 ± 18.3%). All n≥4. **(E)** Representative activation currents of KCNQ1 before (black) and after (red) adding C28 and cAMP. **(F)** Normalized G-V relations of KCNQ1 before (black) and after (red) adding C28 and cAMP. All n≥4.

### ML277 enhances the cAMP sensitivity of native I_Ks_ currents

The β-adrenergic stimulation during physical stresses such as swimming is the main trigger for cardiac events of LQT1 patients, especially in patients whose I_Ks_ lose the cAMP sensitivity, predisposing the patients to high-risk arrhythmias due to compromised cardiac adaptation to stresses (13, 14). The above results suggest that an enhancement of the AO state occupancy can be used as a strategy to increase the response of KCNQ1 to β-adrenergic stimulation in the heart. We next sought to examine this strategy in both native I_Ks_ channels in the heart and mutant-I_Ks_ channels that show defective cAMP sensitivity.

Guinea pig ventricular myocytes provide a good model for studying the function of native I_Ks_ currents due to their relatively high expression level of I_Ks_ channels (33–35). In patch clamp whole-cell mode, we injected a 180 pA/10 ms current pulse to induce action potentials (APs) in guinea pig ventricular myocytes at 1 Hz. 10 µM isoproterenol (ISO, a β-adrenergic agonist) was used to mimic the β-adrenergic stimulation in cardiomyocytes. ISO shortened the APD_90_ and APD_50_, consistent with the repolarization of cardiac APs being accelerated by increasing the I_Ks_ current after PKA-dependent phosphorylation (**Fig. 4A, C, D**). Interestingly, in the presence of 1 µM ML277, the APs were more sensitive to the ISO stimulation, so that ISO induced increased APD_90_ and APD_50_ shortening as compared to the native I_Ks_ without ML277 (**Fig. 4B-D**). Of note, ML277 can only activate KCNQ1 but not I_Ks_ with saturated KCNE1 (21, 36). ML277 could modulate the native I_Ks_ in ventricular myocytes because the native I_Ks_ channels are not fully saturated with KCNE1 association (37–41) (see Discussion). These results suggest that: in the absence of ML277, ISO mainly works on I_Ks_ channels that are associated with KCNE1; ML277 can boost the cAMP sensitivity of native I_Ks_ channels that are not saturated with KCNE1, so that ISO can modify additional native I_Ks_ channels and, therefore, enhance the APD shortening.

**Fig. 4.**
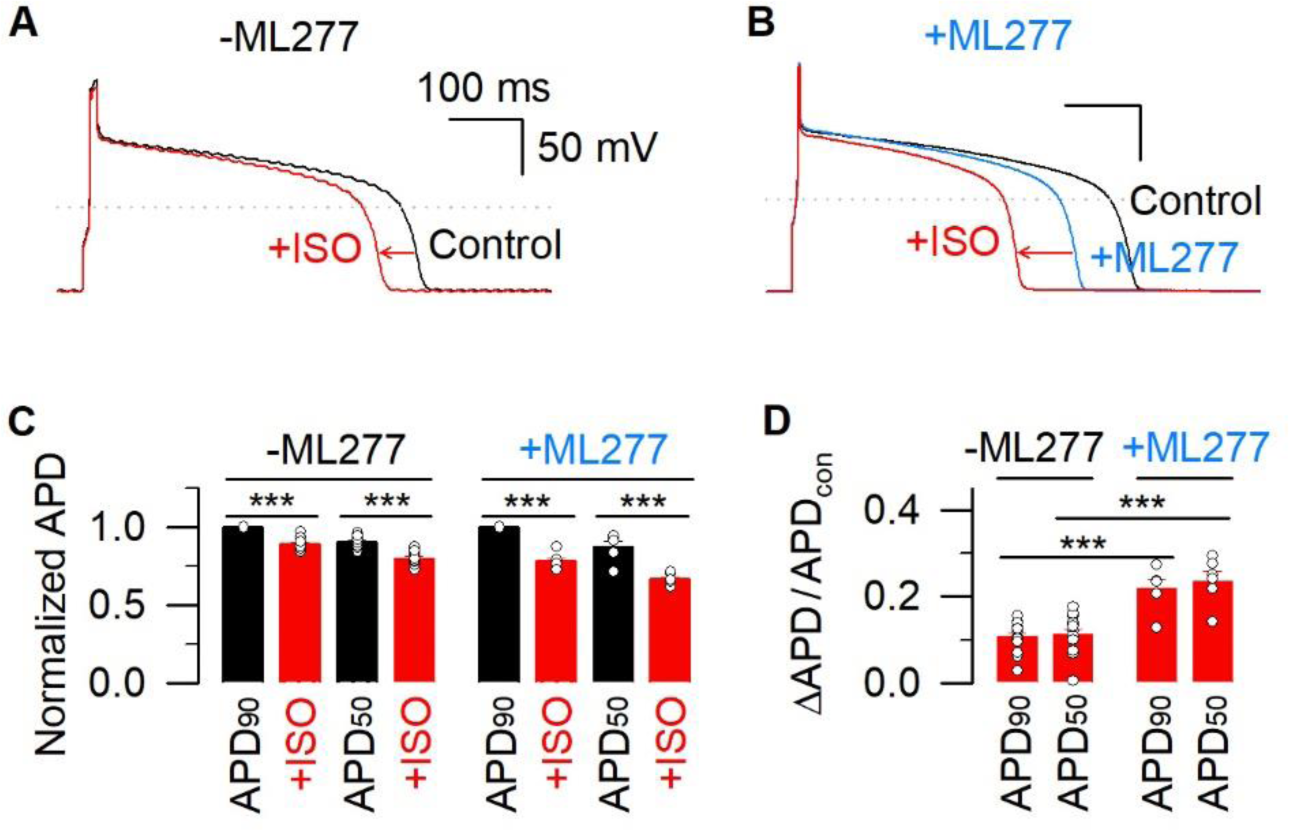
ML277 increases the cAMP sensitivity of native I_Ks_ currents from guinea pig ventricular myocytes. **(A)** Representative action potentials of guinea pig ventricular myocytes before (black) and after (red) adding 10 µM ISO. **(B)** Representative action potentials of guinea pig ventricular myocytes, in the presence of 1 µM ML277, before (blue) and after (red) adding 10 µM ISO. The action potential before adding ML277 from the same cell is shown in black. **(C)** Normalized APD. All APD values are normalized to APD_90_ in the absence of ISO. Black, data in the absence of ISO; red, in the presence of 10 µM ISO. In the absence of ML277, ISO-induced changes were from 1 to 0.89 ± 0.01 for APD_90_ and from 0.91 ± 0.01 to 0.80 ± 0.01 for APD_50_. In the presence of ML277, ISO-induced changes were from 1 to 0.78 ± 0.02 for APD_90_ and from 0.88 ± 0.04 to 0.67 ± 0.02 for APD_50_. All n≥6. **(D)** The ISO-induced changes of APD_90_ and APD_50_. In the absence of ML277, ISO-induced changes were 0.11 ± 0.01 for APD_90_ and 0.11 ± 0.01 for APD_50_. In the presence of ML277, ISO-induced changes were 0.22 ± 0.02 for APD_90_ and 0.24 ± 0.02 for APD_50_. All n≥6.

### ML277 rescues the defective cAMP effects in the high-risk mutation R243C

The KCNQ1 protein sequence contains 676 amino acids. Among these residues more than 300 mutations have been found to associate with LQT1, accounting for ∼35% of all long QT syndrome cases (7, 8). LQT1 is frequently triggered by adrenergic stimuli, and clinical studies found that the reduced cAMP response and the lethality of LQT1 mutations are associated with the location of the mutations in the channel protein (14, 42). For instance, R243C, a “C-loop” mutation located in the S4-S5 linker (**Fig. 5A**), reduces the cAMP effect on the mutant I_Ks_ channels and poses a higher risk of arrhythmia and sudden death to patients than other mutations (14). Rescue of the impaired cAMP sensitivity of R243C patients represents a feasible strategy to alleviate the clinical risk under stressful conditions.

**Fig. 5.**
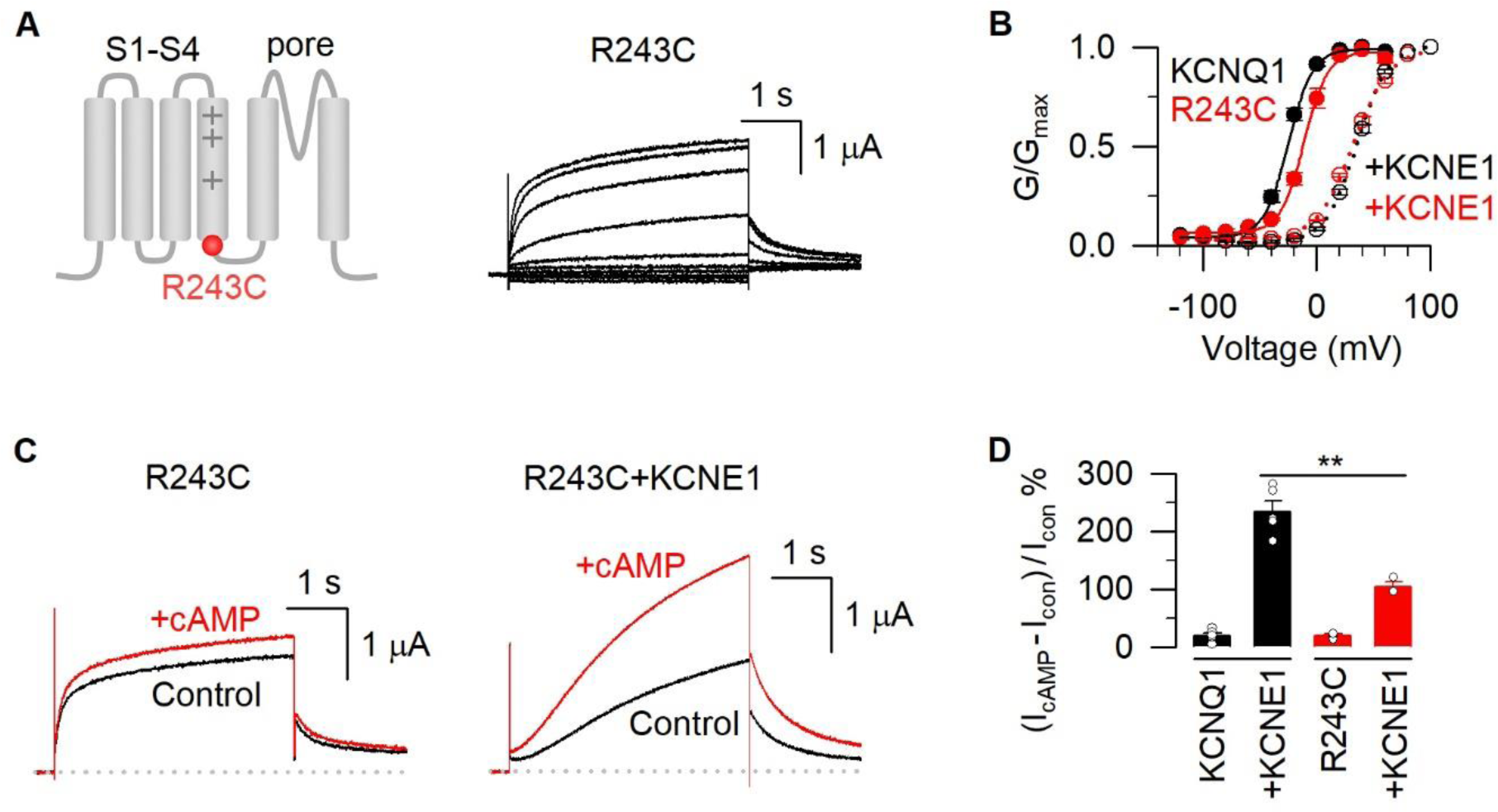
The high risk LQT1 mutation R243C+KCNE1 shows impaired cAMP modulation. **(A)** Left, a cartoon of KCNQ1 to show the location of the high-risk LQT1 mutation R243C in the S4-S5 linker. Right, representative currents of R243C elicited by depolarizing voltage pulses (from −120 to +60 mV). **(B)** G-V relations of WT (black) and R243C KCNQ1 (red), with (solid circles) and without (open circles) KCNE1. All n≥4. **(C)** Representative currents of R243C and R243C+KCNE1 before (black) and after (red) adding 0.5 mM cAMP. **(D)** Average results of cAMP induced current increase from KCNQ1 (19.6 ± 4.8%), KCNQ1+KCNE1 (234.3 ± 18.1%), R243C (21.1 ± 1.9%), and R243C+KCNE1 (104.1 ± 8.0%). All n≥3.

To achieve this goal, we first characterized the voltage-dependent activation of R243C and R243C-I_Ks_ channels. We found that these channels have similar G–V relations as the WT KCNQ1 and I_Ks_ channels (**Fig. 5B**). Western blot data in previous studies showed that R243C has a robust membrane expression similar to the WT KCNQ1 (14). However, R243C significantly suppresses KCNE1’s effect to boost cAMP sensitivity: the mutant R243C itself shows a low cAMP modulation (21.1 ± 1.9%, **Fig. 5C, D**), and R243C+KCNE1 also loses most of the cAMP sensitivity as compared to the WT I_Ks_ channel (104.1 ± 8.0%, **Fig. 5C, D**). These properties suggest that the defective cAMP modulation of R243C-I_Ks_ makes a major contribution to the increased clinical risk of the patients carrying this mutation under stress conditions (14).

Our results in **Fig. 1** showed that the KCNE1 subunit boosts the cAMP sensitivity of the I_Ks_ channel by enhancing the AO state of KCNQ1. In the high-risk LQT1 mutation R243C when the KCNE1’s boosting effect is largely reduced, we utilized ML277 as an alternative means to enhance the AO state occupancy of R243C and measured the cAMP effects. Similar to the WT KCNQ1 channel, 1 µM ML277 enhanced the current amplitude by increasing the AO state occupancy (**Fig. 6A**). We then applied cAMP and measured the current amplitude increase. ML277 clearly boosted the cAMP sensitivity, and thus the cAMP induced current amplitude more than that of R243C itself (**Fig. 6A, B**). Furthermore, the G-V relation of R243C after adding ML277+cAMP also shows a ∼9 mV shift to more positive voltages (**Fig. 6C**). Interestingly, compared to the R243C channel that show minimum cAMP effect, the same concentration of cAMP induced a greater current increase for R243C+1 µM ML277 (75.7 ± 7.3%, **Fig. 6D, E**).

**Fig. 6.**
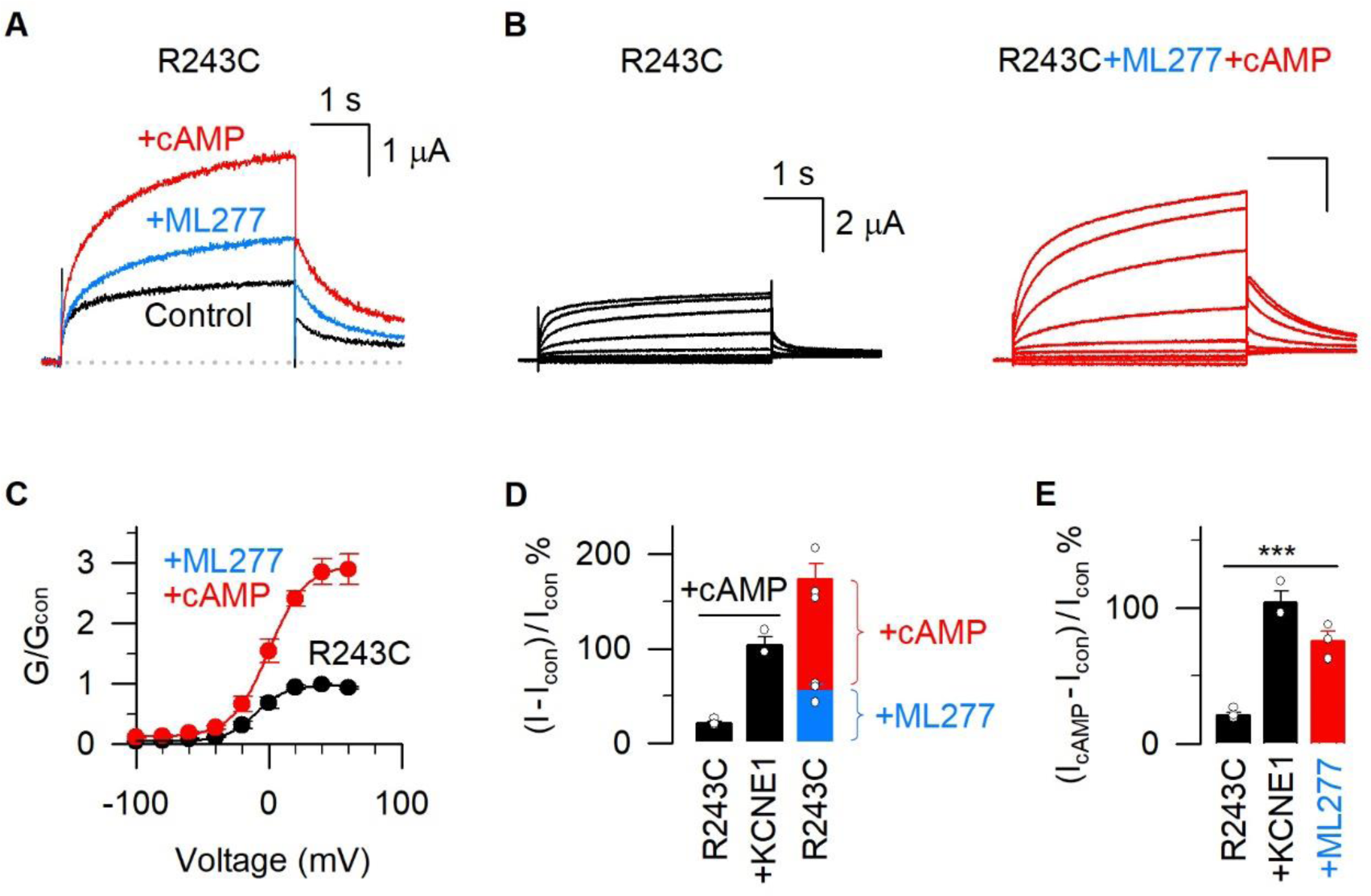
ML277 rescues the defective cAMP effects in the high-risk mutation R243C. **(A)** Currents of R243C channel before (black) and after (blue) adding ML277, and then after adding cAMP (red). **(B)** Representative activation currents of R243C before (black) and after (red) adding ML277 and cAMP. **(C)** G-V relations of R243C before (black) and after (red) adding ML277 and cAMP. Data are normalized to control. **(D)** Current increases of R243C after adding ML277 (blue, 55.6 ± 6.2%) and cAMP (red, 173.5 ± 16.7%). The cAMP-induced current increases of R243C (21.1 ± 1.9%) and R243C+KCNE1 (104.1 ± 8.0%) channels were shown in black. All n≥4. **(E)** The cAMP-induced current increases of R243C (21.1 ± 1.9%) and R243C+KCNE1 (104.1 ± 8.0%), and R243C after adding ML277 (75.7 ± 7.3%). The data showed significant differences (“***”) by using Anova analysis. All n≥3.

## Discussion

In this study, we leverage the unique two open states gating mechanisms of the KCNQ1 channel and the state-dependent mutant channels that selectively open into the IO (S338F and E1R/R2E) and AO (F351A and E1R/R4E) states and find that the AO state is mainly responsible for the cAMP sensitivity of the KCNQ1 and I_Ks_ channels (**Fig. 1**). This finding answers the long-standing question why the KCNQ1 channel shows only a small current increase, while the I_Ks_ channel shows a large current increase under the β-adrenergic stimulation: KCNQ1 predominantly opens in the IO state so that it shows mainly the less cAMP sensitive IO state phenotype; the KCNE1 association, on the other hand, suppresses the IO state and enhances the AO state and therefore boosts the cAMP sensitivity of KCNQ1. Taking advantage of the unique feature that small molecules ML277 and C28 can specifically enhance the AO state, we found that they effectively enhances the cAMP sensitivity of the KCNQ1 channel (**Figs. 2,3**). This finding suggests that the cAMP sensitivity of the KCNQ1 channel originates from the AO state of the channel itself, independent of the KCNE1 association, and increasing the AO state occupancy in KCNQ1 may enhance its cAMP sensitivity. These results suggest that enhancing the AO state occupancy of KCNQ1 channels by exogenous modulators can be a new strategy to increase the cAMP sensitivity of native I_Ks_ channels in the heart. We confirmed this strategy on native I_Ks_ channels in guinea pig ventricular myocytes and the high-risk LQT1 mutation R243C. ML277 effectively sensitized the cAMP responses of APs from guinea pig ventricular myocytes (**Fig. 4**) and the high-risk LQT1 mutation R243C (**Figs. 5,6**).

The cAMP-dependent modulation of the I_Ks_ channel plays a key role in the cardiac adaptation under stress conditions. For LQT1 patients, β-adrenergic stimulation is the main trigger for cardiac events. LQT1 mutations that reduce cAMP sensitivity of the mutant-I_Ks_ currents show higher risk of cardiac events and increased lethality (13, 14). However, so far, no strategy has been reported to rescue this defective cAMP modulation. Our findings in this study clearly show that enhancing the AO state occupancy of KCNQ1 channels can be a new anti-arrhythmic strategy, especially for the high-risk variants where the KCNE1’s effect of boosting cAMP sensitivity is largely suppressed.

Interestingly, several lines of evidence have demonstrated that, in native I_Ks_ currents, KCNQ1 can associate with KCNE1 at various stoichiometries. First, *in vitro* stoichiometry studies show that one tetrameric KCNQ1 channel can associate with different numbers of KCNE1 (from 0 to 4), depending on the KCNQ1:KCNE1 expression ratio (37–39, 41, 43–45). Second, *in vivo* studies in ventricular myocytes suggest that the ratio of KCNQ1:KCNE1 does not remain stable but changes dynamically in different conditions over time and pathology progression (40, 46–48). Third, the G–V relations of native I_Ks_ currents from neonatal mouse, guinea pig, and rabbit ventricular myocytes fall in between that of the KCNQ1 only and I_Ks_ with saturated KCNE1 association (34, 49). Fourth, although ML277 selectively activates the KCNQ1 channel but not the I_Ks_ channel with saturated KCNE1 association, pharmacological studies in guinea pig ventricular myocytes, human induced pluripotent stem cell (iPSC) derived cardiomyocytes, and zebrafish hearts show that ML277 effectively shortens the APD via activating native I_Ks_ currents (**Fig. 4**) (36, 50, 51). These results consistently suggest that, in cardiomyocytes the native I_Ks_ channels are not fully saturated with KCNE1 association. The KCNQ1 subunits not associated with KCNE1 constitute the “KCNQ1 component” in native I_Ks_ channels that can be further modulated by small molecules such as ML277. Consistent with this finding, the Fedida lab has quantified the cAMP effects on I_Ks_ channels with different KCNQ1:KCNE1 stoichiometry, and they found that cAMP induces no V_50_ shift on the WT KCNQ1 channel, but gradually increases with the number of KCNE1 binding to the I_Ks_ channel (32). These results indicate that our proposed anti-arrhythmic strategy of boosting cAMP sensitivity of native and mutant I_Ks_ by increasing the AO state of the channel using exogenous modulators can be effective in treating LQT1 patients.

## Materials and Methods

### Constructs and mutagenesis

Overlap extension and high-fidelity PCR were used for making KCNQ1 channel point mutations. Each KCNQ1 mutation was verified by DNA sequencing. Then cRNA of WT and mutant KCNQ1 were synthesized using the mMessage T7 polymerase kit (Applied Biosystems-Thermo Fisher Scientific) for oocyte injections.

### Oocyte preparation and channel expression

Oocytes (at stage V or VI) were obtained from *Xenopus laevis* by laparotomy. All procedures are consistent with the recommendations of the Panel on Euthanasia of the American Veterinary Medical Association. Oocytes were digested by collagenase (0.5 mg/ml, Sigma Aldrich) and micro-injected with KCNQ1 and KCNE1 cRNAs. WT or mutant KCNQ1 cRNAs (9.2 ng) with or without KCNE1 cRNA were injected into each oocyte with a KCNQ1:KCNE1 weight ratio of 4:1, which could saturate the KCNE1 association to KCNQ1 (21, 38). Injected cells were kept in ND96 solution (in mM): 96 NaCl, 2 KCl, 1.8 CaCl_2_, 1 MgCl_2_, 5 HEPES, 2.5 CH_3_COCO_2_Na, 1:100 Pen-Strep, pH 7.6) at 18°C for 2-6 days for electrophysiology recordings.

### Two-electrode voltage clamp (TEVC)

Microelectrodes (Sutter Instrument, Item #: B150-117-10) were made with a Sutter (P-97) puller with 1-3 MΩ resistances when filled with 3 M KCl. The extracellular solution was ND96 solution without CH_3_COCO_2_Na. Currents were recorded with a CA-1B amplifier (Dagan, Minneapolis, MN) with Patchmaster (HEKA) software. Signals were sampled at 1 kHz and low-pass-filtered at 2 kHz. ML277 and cAMP stocks (Sigma Aldrich) were added to the bath and diluted to working concentrations. All recordings were performed at room temperature (21–23 °C). Data were analyzed with Clampfit (Axon Instruments, Inc., Sunnyvale, CA), Sigmaplot (SPSS, Inc., San Jose, CA), and IGOR (Wavemetrics, Lake Oswego, OR).

### Endogenous Yotiao detection by RT-PCR

The A-kinase anchoring protein (AKAP, also named Yotiao) is required for the PKA-dependent phosphorylation of the I_Ks_ channel (9). We found that, in *Xenopus* oocytes, the I_Ks_ channel without human Yotiao (hYotiao) injection is activated by cAMP to a similar extent to that after hYotiao injection (KCNQ1:hYotiao mRNA weight ratio was 1:1), which suggests that there is endogenous frog Yotiao (fYotiao) in *Xenopus* oocytes that has similar function to the hYotiao (**Fig. S2**).

To confirm the existence of endogenous fYotiao in *Xenopus* oocytes, total RNAs were extracted from 30 hYotiao injected or uninjected *Xenopus* Oocytes. Experiments were carried out using a Purelink RNA extract mini kit (Thermo Fisher Scientific, USA). Subsequently, the isolated RNA was converted to cDNA using an iScript cDNA Synthesis Kit (Bio-Rad, USA). The polymerase chain reaction (PCR) was conducted using Herculase II Fusion DNA Polymerase (Agilent, USA). The sequences of both f- and h-Yotiao were acquired from the National Center for Biotechnology Information (NCBI). The oligonucleotide forward primer sequence used for both f- and h-Yotiao is GAGCAGCTGAGTTCTGAGA. The reverse primer sequences were GGCCGAGACTGTCCAGTA for h-Yotiao, and CTGTGTGAATCAGACTGTTC for f-Yotiao. The amplified DNA fragments for the human F-R primer pair have a length of 557 base pairs (bp), while the amplified DNA fragments for the frog F-R primer pair have a length of 563 bp (**Fig. S2**). The endogenous fYotiao protein was found to have a similar function to the hYotiao in the PKA-dependent phosphorylation of the I_Ks_ channel (**Fig. S2**). Therefore, we did not inject hYotiao in other experiments with oocytes.

### Guinea pig cardiomyocytes preparation and action potential recording

Cardiac myocyte isolation: The use of guinea pigs was approved by the Stony Brook University Institutional Animal Care and Use Committee. Single ventricular myocytes were acutely enzymatically isolated from guinea pig heart as previously described(52). Guinea pigs, weighing 300-500 g, were sacrificed by peritoneal injection of sodium pentobarbitone (1 ml, 390mg/ml). The isolated cells were stored in KB solution containing (in mM) 83 KCl, 30 K_2_HPO_4_, 5 MgSO_4_, 5 Na-pyruvate, 5 β-OH-butyric acid, 20 creatine, 20 taurine, 10 glucose, 0.5 EGTA, pH 7.2. Electrophysiological recording of action potentials: Action potentials were recorded with whole cell current clamp recording. Freshly isolated guinea pig cardiac myocytes were paced at 1 Hz with a 180 pA pulse current for 10 ms duration to generate action potentials. The APD_90_ and APD_50_ were determined at 90% and 50% repolarization from the peak amplitude, respectively. Several minutes were allowed for the APD to reach steady state before data was collected. The extracellular solution contains (in mM): 140 NaCl, 3 KCl, 1 MgCl_2_, 1.8 CaCl_2_, 10 HEPES, 10 Glucose, pH 7.4. The pipette solution contains (in mM): 115 K-Aspartic Acid, 35 KOH, 3 MgCl_2_, 10 HEPES, 11 EGTA, 5 Glucose, 3 MgATP, pH 7.4. All recordings were performed at room temperature (21–23 °C).

## Acknowledgments

This work was supported by NIH grants RO1 HL126774, HL155398 and US-Israel Binational Science Foundation research grant 2019159 to J.C.; HL166628 to I.S.C. and AHA postdoctoral fellowship 18POST34030203 to P.H.. P.H. was also supported by the following grants in writing the manuscript: National Natural Science Foundation of China (32171221) and Macau Science and Technology Development Fund (0074/2022/A2 and 0098/2023/RIA2).

## Author Contributions

P.H. and J.C. conceived this work. P.H., L.Z., L.Zhong., J.S., H.W., J.G., H.C., and J.Z. performed the experiments. P.H., I.S.C. and J.C. analyzed data. P.H. and J.C. wrote the paper with input from all authors.

## Data Availability

All study data are included in the article and/or supporting information.

## Supplementary Figures

**Figure S1.**
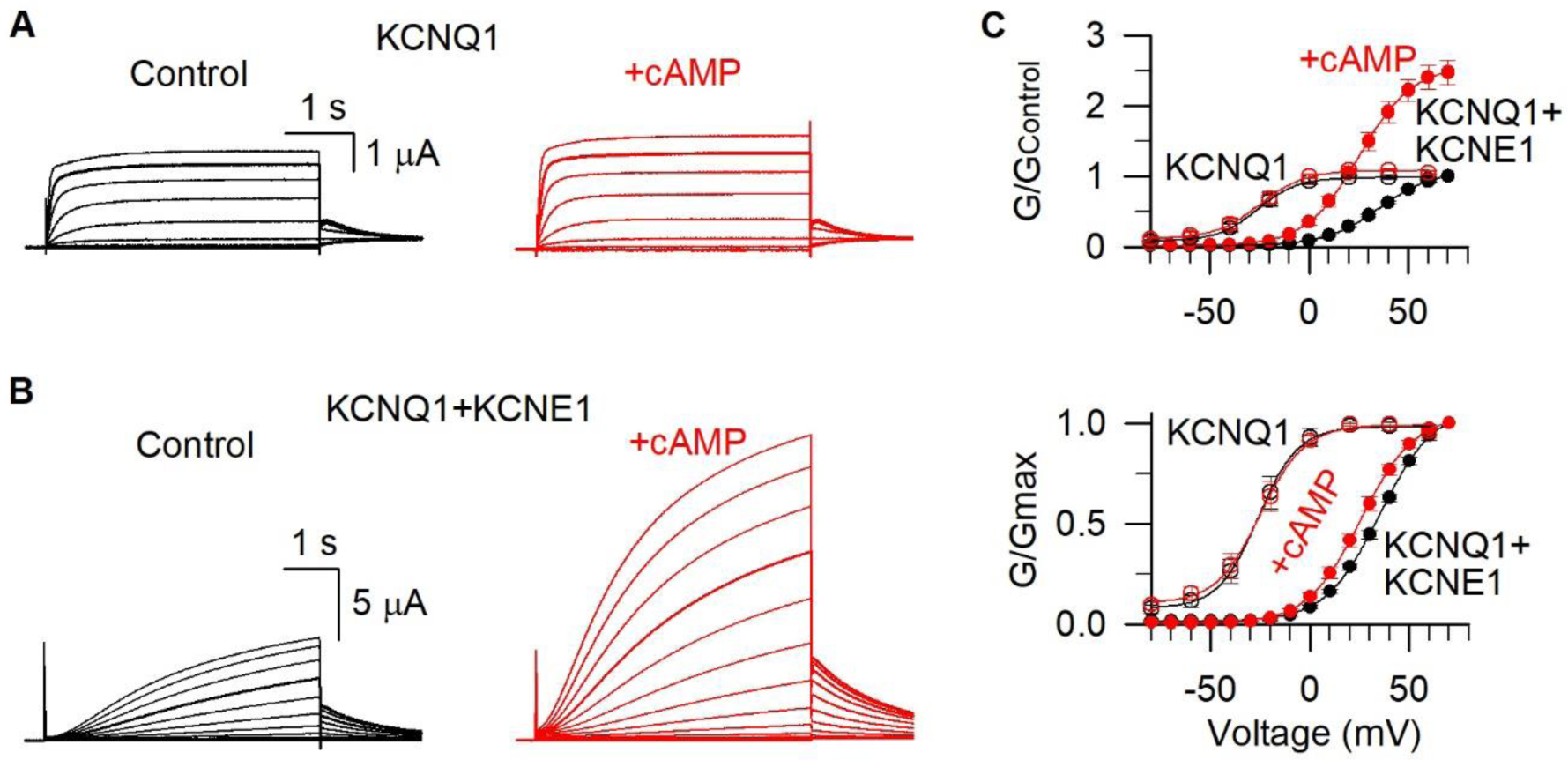
The KCNE1 subunit is required for the cAMP-induced current increase. **(A-B)** Representative currents of KCNQ1 and KCNQ1+KCNE1 channels before (black) and after (red) adding 0.5 mM cAMP. The test pulse was from −100 to +60/+70 mV for 4 s and then returned to −40 mV for recording tail currents. Currents recorded at +40 mV were highlighted. **(C)** Top, conductance-voltage (G–V) relations of KCNQ1 (open circles) and KCNQ1+KCNE1 (solid circles) before (black) and after (red) adding cAMP were normalized to control (KCNQ1 without cAMP). Bottom, normalized G–V relations of KCNQ1 (open circles) and KCNQ1+KCNE1 (solid circles) channels before (black) and after (red) adding cAMP. The V_50_s were −25.0 ± 3.7 mV (control) and −26.1 ± 3.3 mV (adding cAMP) for KCNQ1, and 35.4 ± 1.4 mV (control) and 25.8 ± 2.0 mV (adding cAMP) for KCNQ1+KCNE1. All n≥4.

**Figure S2.**
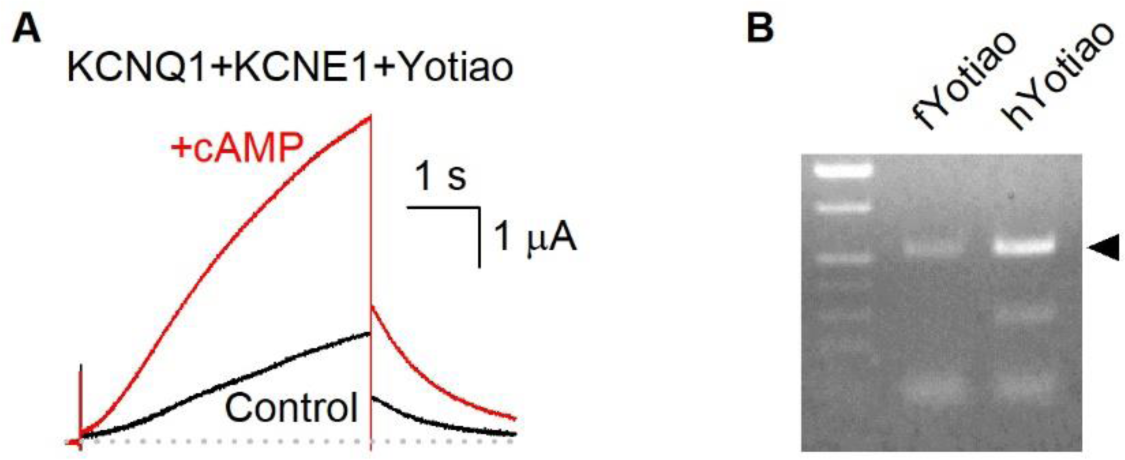
The endogenous Yotiao was detected in *Xenopus* oocytes. **(A)** Representative currents of KCNQ1+KCNE1+hYotiao before (black) and after (red) adding 0.5 mM cAMP. The test pulse was +40 mV for 4 s. The cAMP induced I_Ks_ current increase is comparable with that in the absence of hYotiao injection (**Figure 1**). n≥3. **(B)** cDNA detection of endogenous Yotiao (fYotiao) in *Xenopus* oocytes. Human Yotiao (hYotiao) detected in hYotiao injected oocytes is shown as positive control.

