## SUPPLEMENTARY MATERIALS for "The fully activated open state of KCNQ1 controls the cardiac “fight-or-flight” response"

^&:^ Send correspondence to;

**Supplementary figures**

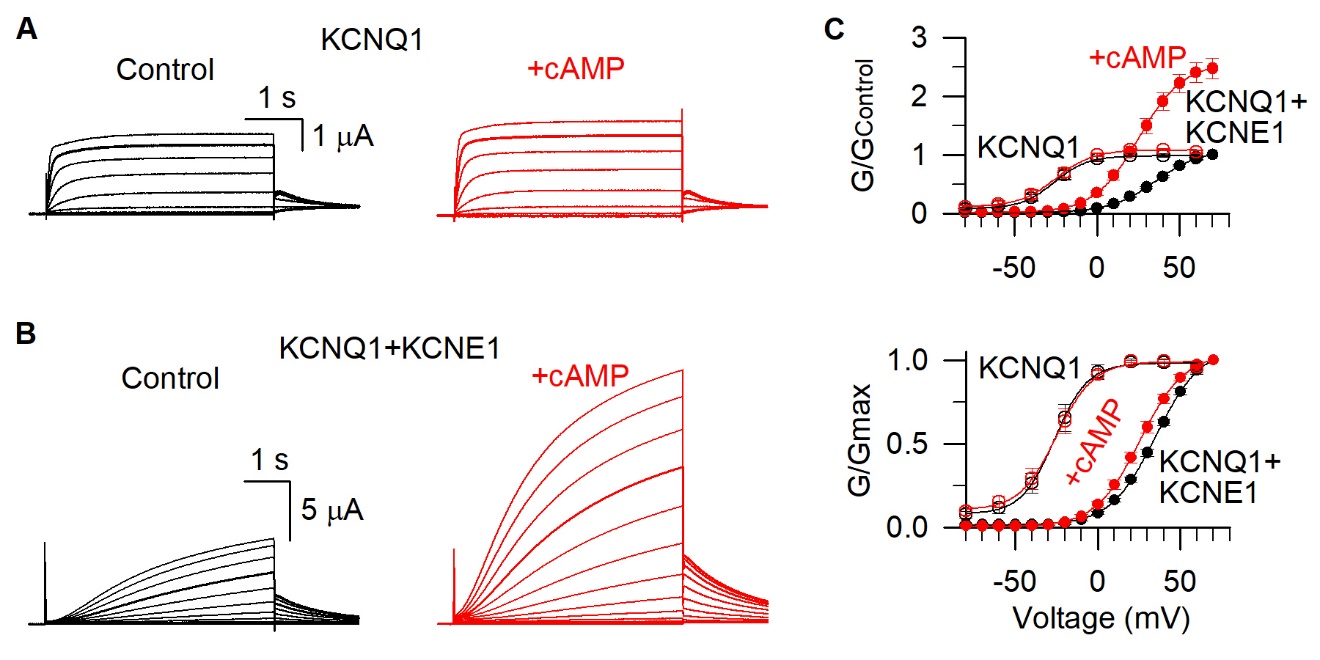

**Figure S1. The KCNE1 subunit is required for the cAMP-induced current increase. (A-B)** Representative currents of KCNQ1 and KCNQ1+KCNE1 channels before (black) and after (red) adding 0.5 mM cAMP. The test pulse was from -100 to +60/+70 mV for 4 s and then returned to -40 mV for recording tail currents. Currents recorded at +40 mV were highlighted. **(C)** Top, conductance-voltage (G–V) relations of KCNQ1 (open circles) and KCNQ1+KCNE1 (solid circles) before (black) and after (red) adding cAMP were normalized to control (KCNQ1 without cAMP). Bottom, normalized G–V relations of KCNQ1 (open circles) and KCNQ1+KCNE1 (solid circles) channels before (black) and after (red) adding cAMP. The V_50_s were -25.0 ± 3.7 mV (control) and -26.1 ± 3.3 mV (adding cAMP) for KCNQ1, and 35.4 ± 1.4 mV (control) and 25.8 ± 2.0 mV (adding cAMP) for KCNQ1+KCNE1. All n≥4.

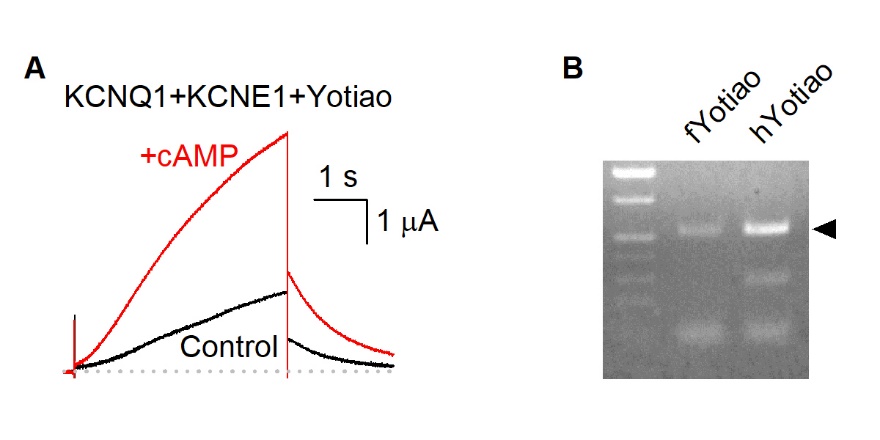

**Figure S2. The endogenous Yotiao was detected in *Xenopus* oocytes. (A)** Representative currents of KCNQ1+KCNE1+hYotiao before (black) and after (red) adding 0.5 mM cAMP. The test pulse was +40 mV for 4 s. The cAMP induced I_Ks_ current increase is comparable with that in the absence of hYotiao injection (**Figure 1**). n≥3. **(B)** cDNA detection of endogenous Yotiao (fYotiao) in *Xenopus* oocytes. Human Yotiao (hYotiao) detected in hYotiao injected oocytes is shown as positive control.
